# Visualization of SARS-CoV-2 infection dynamic

**DOI:** 10.1101/2021.06.03.446942

**Authors:** Chengjin Ye, Kevin Chiem, Jun-Gyu Park, Jesus A. Silvas, Desarey Morales Vasquez, Jordi B. Torrelles, James J. Kobie, Mark R. Walter, Juan Carlos de la Torre, Luis Martinez-Sobrido

## Abstract

Replication-competent recombinant viruses expressing reporter genes provide valuable tools to investigate viral infection. Low levels of reporter gene expressed from previous reporter-expressing rSARS-CoV-2 have jeopardized their use to monitor the dynamics of SARS-CoV-2 infection *in vitro* or *in vivo*. Here, we report an alternative strategy where reporter genes were placed upstream of the viral nucleocapsid gene followed by a 2A cleavage peptide. The higher levels of reporter expression using this strategy resulted in efficient visualization of rSARS-CoV-2 in infected cultured cells and K18 hACE2 transgenic mice. Importantly, real-time viral infection was readily tracked using a non-invasive *in vivo* imaging system and allowed us to rapidly identify antibodies which are able to neutralize SARS-CoV-2 infection *in vivo*. Notably, these reporter-expressing rSARS-CoV-2 retained wild-type virus like pathogenicity *in vivo*, supporting their use to investigate viral infection, dissemination, pathogenesis and therapeutic interventions for the treatment of SARS-CoV-2 *in vivo*.

## INTRODUCTION

Coronaviruses (CoVs) are enveloped, single-stranded, positive-sense RNA viruses that belong to the *Coronaviridae* family that can cause mild to severe respiratory infections in humans (Hu et al., 2020). Two CoVs have been associated with severe respiratory syndrome in the past two decades: Severe Acute Respiratory Syndrome CoV (SARS-CoV) in 2002-2003 and Middle East Respiratory Syndrome CoV (MERS-CoV) in 2012-present (De Wit et al., 2016). Severe acute respiratory syndrome coronavirus 2 (SARS-CoV-2) emerged in the Chinese city of Wuhan in December 2019 and is the causative agent of the coronavirus disease 2019 (COVID-19) pandemic (Lu et al., 2020; Wu et al., 2020). As of June 2021, SARS-CoV-2 has been reported to be responsible for over 150 million human infection cases and more than 3 million deaths around the World (https://covid19.who.int/).

Like SARS-CoV and MERS-CoV, SARS-CoV-2 mainly replicates in the upper (nasal turbinate) and lower (lungs) respiratory tract, resulting, in some cases, in fatal respiratory illness (Li et al., 2020; Xu et al., 2020). However, the intra-host dissemination and pathogenesis of SARS-CoV-2 are not well understood. Several animal models of SARS-CoV-2 infection have been established and have already provided very valuable information to understand the mechanism of tissue and cell tropism, replication and pathogenesis (Imai et al., 2020; Kim et al., 2020; Oladunni et al., 2020; Singh et al., 2021; Sun et al., 2020). However, assessing the presence of SARS-CoV-2 in infected animals, organs or tissues has required collection and processing of samples upon euthanasia, which complicates studies examining the longitudinal dynamic of a viral infection within an infected host. Recombinant (r)SARS-CoV-2 expressing reporter genes could overcome this problem and allow tracking viral infection *in vivo* and in real time by monitoring the expression of the reporter gene. We and others have documented the feasibility of generating reporter-expressing rSARS-CoV-2 using reverse genetic system (Chiem et al., 2021; Xie et al., 2020a). These rSARS-CoV-2 have been genetically engineered to express the reporter gene by substituting the viral open reading frame (ORF) 7a protein with the reporter gene of interest, an experimental approach first employed to generate reporter-expressing rSARS-CoV (Sims et al., 2005). Despite these reporter-expressing rSARS-CoV-2 were showing comparable plaque phenotype, replication and growth kinetics as those of wild-type virus (rSARS-CoV-2/WT) *in vitro* (Chiem et al., 2021; Xie et al., 2020a; Xie et al., 2020b), it is unclear if the reporter-expressing rSARS-CoV-2 lacking ORF7a recapitulate viral pathogenicity *in vivo* and whether reporter gene expression levels could be efficiently tracked *ex vivo* using tissues or organs from infected animals, or in a whole organism *in vivo*.

In this study, we cloned fluorescent (Venus) and luciferase (Nano luciferase, Nluc) reporter genes upstream of the SARS-CoV-2 nucleocapsid (N) gene separated by the porcine tescherovirus (PTV-1) 2A proteolytic cleavage site to generate new reporter-expressing rSARS-CoV-2 without the deletion of the ORF7a protein. *In vitro*, rSARS-CoV-2 expressing reporter genes from the viral N locus replicated and made viral plaques similar to those of rSARS-CoV-2/WT. Notably, reporter-expressing rSARS-CoV-2 generated using this 2A strategy expressed higher levels of reporter gene expression compared to those rSARS-CoV-2 generated by substituting the viral ORF7a protein with the reporter gene of interest. Importantly, rSARS-CoV-2/Venus-2A and rSARS-CoV-2/Nluc-2A showed rSARS-CoV-2/WT-like pathogenicity *in vivo*. Notably, the higher level of Venus expression from rSARS-CoV-2/Venus-2A allowed us to detect viral infection in the lungs of infected K18 human angiotensin converting enzyme 2 (hACE2) transgenic mice using an *in vivo* imaging system (IVIS). Moreover, Venus expression from rSARS-CoV-2/Venus-2A was stable up to seven passages *in vitro* in cultured Vero E6 cells and *in vivo* up to day 6 post-infection. Importantly, levels of Venus expression correlated well with viral titers detected in the lungs, demonstrating the feasibility of using Venus expression as a valid surrogate marker to evaluate SARS-CoV-2 infection. Using rSARS-CoV-2/Nluc-2A, we were able to track the dynamics of viral infection in real time and longitudinally assess SARS-CoV-2 infection *in vivo*. Finally, we testified the feasibility of using the rSARS-CoV-2/Nluc-2A to rapidly and accurately identify antibodies that neutralize viral infection *in vivo*.

Our data demonstrate that these next-generation of rSARS-CoV-2 expressing reporter genes we have generated can be used to easily monitor viral infection in cultured cells and in validated animal models of infection. Importantly, our new rSARS-CoV-2/Venus-2A or rSARS-CoV-2/Nluc-2A retain similar virulence to that of rSARS-CoV-2/WT in K18 hACE2 transgenic mice and can be used to investigate viral replication, tropism and viral dissemination and pathogenesis *in vivo* and to rapidly identify therapeutics for the treatment of SARS-CoV-2 infection and associated COVID-19 disease.

## RESULTS

### Generation of rSARS-CoV-2 expressing Venus

We have recently described the generation and characterization of rSARS-CoV-2 where a reporter gene of interest replaced the viral ORF7a protein (Chiem et al., 2021). However, these rSARS-CoV-2 showed low levels of reporter gene expression during viral infection. To increase expression levels of reporter gene during SARS-CoV-2 infection and avoid the deletion of ORF7a protein, we implemented a strategy we previously used to generate recombinant influenza viruses and mammarenaviruses (Nogales et al., 2019; Ye et al., 2020b). In this approach, the sequences of the reporter Venus fluorescent protein and the porcine teschovirus 1 (PTV-1) 2A self-cleaving peptide (Caì et al., 2018) were cloned upstream of the SARS-CoV-2 N gene (**Fig. 1A**), and the sequences harboring the fusion Venus-2A-N were cloned into the previously described bacterial artificial chromosome (BAC) containing the entire SARS-CoV-2 genome (Chiem et al., 2020; Ye et al., 2020a). We rescued rSARS-CoV-2/Venus-2A virus according to our previously described protocol (Chiem et al., 2020; Ye et al., 2020a). Vero E6 cells infected with the tissue culture supernatant from Vero E6 cells transfected with the BAC containing the viral genome of rSARS-CoV-2/Venus-2A resulted in the expression of Venus in the same cells expressing the viral N protein (**Fig. 1B**). Venus was readily detected in whole cell lysate from rSARS-CoV-2/Venus-2A-, but not rSARS-CoV-2/WT-infected Vero E6 cells, while the viral N protein was detected in lysate obtained from both rSARS-CoV-2/Venus-2A- and rSARS-CoV-2/WT-infected Vero E6 cells (**Fig. 1C**). We confirmed the genetic identity of rSARS-CoV-2/Venus-2A using reverse-transcription polymerase chain reaction (RT-PCR) to amplify the Venus sequence and the entire sequence between ORF8 and N. The Venus fragment was amplified from cells infected with rSARS-CoV-2/Venus-2A, while the fragment between ORF8 and N was detected in cells infected with either rSARS-CoV-2/WT or rSARS-CoV-2/Venus-2A. As predicted, the amplified fragment from rSARS-CoV-2/Venus-2A infected cells had a higher molecular size than the one obtained from rSARS-CoV-2/WT-infected cells (**Fig. 1D**).

**Figure 1.**
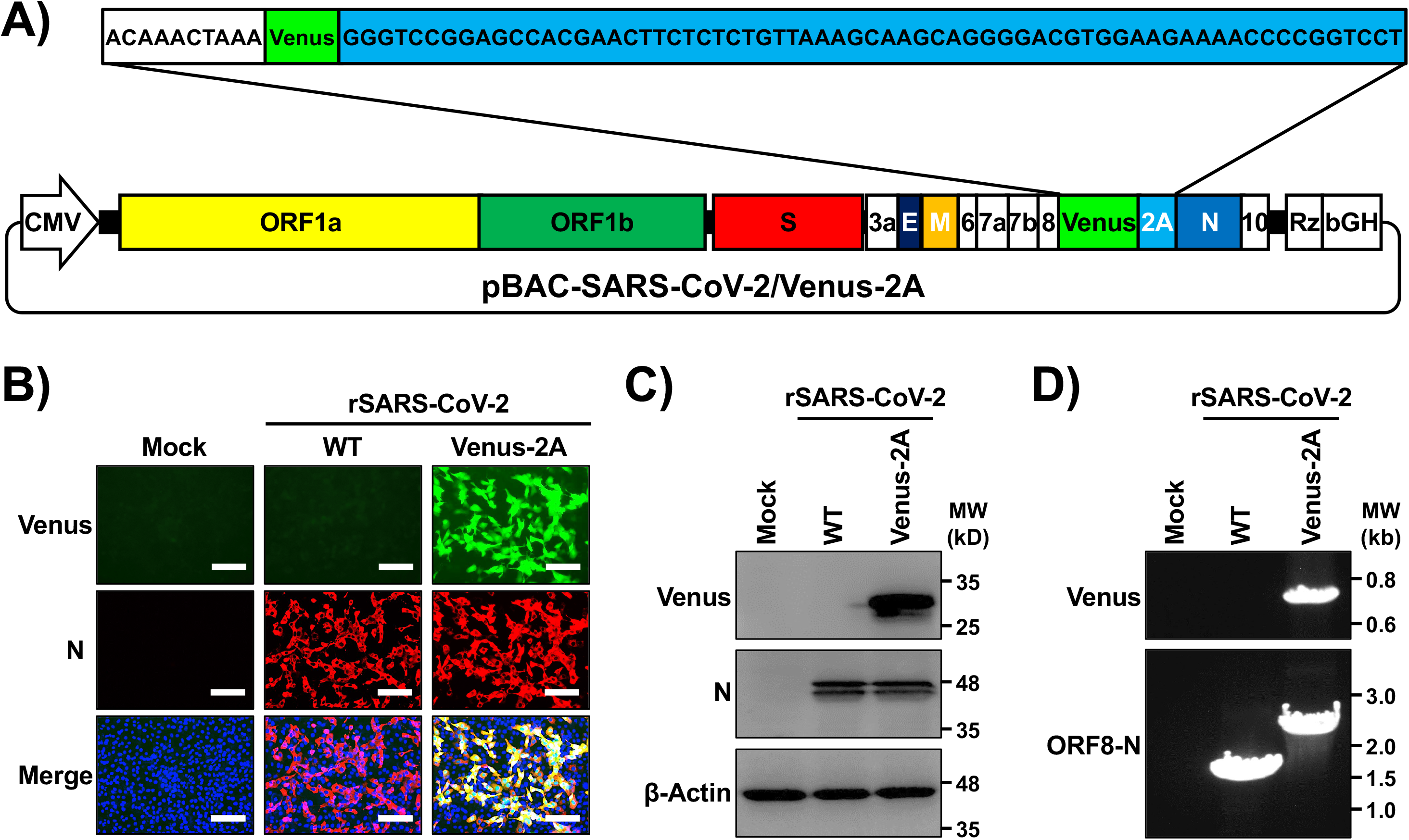
Generation of a rSARS-CoV-2 expressing Venus-2A. **(A)** Schematic representation of the BAC for generation of rSARS-CoV-2/Venus-2A. The sequence encoding the fusion construct Venus-2A was inserted in the viral genome of SARS-CoV-2 in the BAC. The white box represents the intergenic region between ORF8 and N. The green box represents Venus. PTV-1 2A is indicated in light blue. **(B)** Vero E6 cells were mock-infected or infected with rBAC-SARS-CoV-2/WT or rSARS-CoV-2/Venus-2A for 48 h, fixed and immunostained with a MAb against the viral N protein (1C7C7). Cell nuclei were stained with DAPI. Representative images are shown. Scale bars, 100 μm. **(C)** Whole cell lysates from Vero E6 cells mock-infected or infected with rSARS-CoV-2 WT or Venus-2A for 48 h were subjected to Western blot analysis using antibodies against Venus and the viral N protein (1C7C7). β-actin was used as a loading control. **(D)** Total cellular RNA from Vero E6 cells mock-infected or infected with WT or Venus-2A rSARS-CoV-2 was isolated at 48 hpi. RT-PCR was used to amplify Venus (top) or the region between the ORF8 and N proteins (bottom), and the products were separated on a 0.7% agarose gel.

### *In vitro* characterization of rSARS-CoV-2/Venus-2A

We next assessed the fitness of rSARS-CoV-2/Venus-2A in Vero E6 cells by evaluating its growth kinetics, plaque phenotype, and reporter expression and compared them to our previously described rSARS-CoV-2/Δ7a-Venus and rSARS-CoV-2/WT. rSARS-CoV-2/Venus-2A showed similar growth kinetics to those of rSARS-CoV-2/WT or rSARS-CoV-2/Δ7a-Venus (**Fig. 2A**). Likewise, rSARS-CoV-2/Venus-2A, rSARS-CoV-2/WT and rSARS-CoV-2/Δ7a-Venus exhibited similar plaque phenotypes (**Fig. 2B**). Notably, plaques formed by rSARS-CoV-2/Venus-2A, but not those of rSARS-CoV-2/Δ7a-Venus, could be readily detected using a fluorescent imaging system, most likely because of the higher levels of Venus expressed from the locus of the viral N than those from the locus of the viral ORF7a (**Fig. 2B**). We next examined Venus expression in Vero E6 cells infected (MOI 0.001) with either rSARS-CoV-2/Venus-2A or rSARS-CoV-2/Δ7a-Venus. Both infections showed peak Venus expression at 48 h post-infection (hpi), but Venus expression levels from rSARS-CoV-2/Venus-2A infected Vero E6 cells were higher than those infected with rSARS-CoV-2/Δ7a-Venus (**Fig. 2C**). To confirm this, we prepared whole cell lysates from Mock, rSARS-CoV-2/Venus-2A and rSARS-CoV-2/Δ7a-Venus infected Vero E6 cells and analyzed them by Western blot. Venus expression levels in rSARS-CoV-2/Venus-2A infected Vero E6 cells were greater than those of rSARS-CoV-2/Δ7a-Venus infected cells at all time points, whereas expression levels of N protein were comparable at all time points in Vero E6 cells infected with rSARS-CoV-2/Venus-2A and rSARS-CoV-2/Δ7a-Venus (**Fig. 2D**).

**Figure 2.**
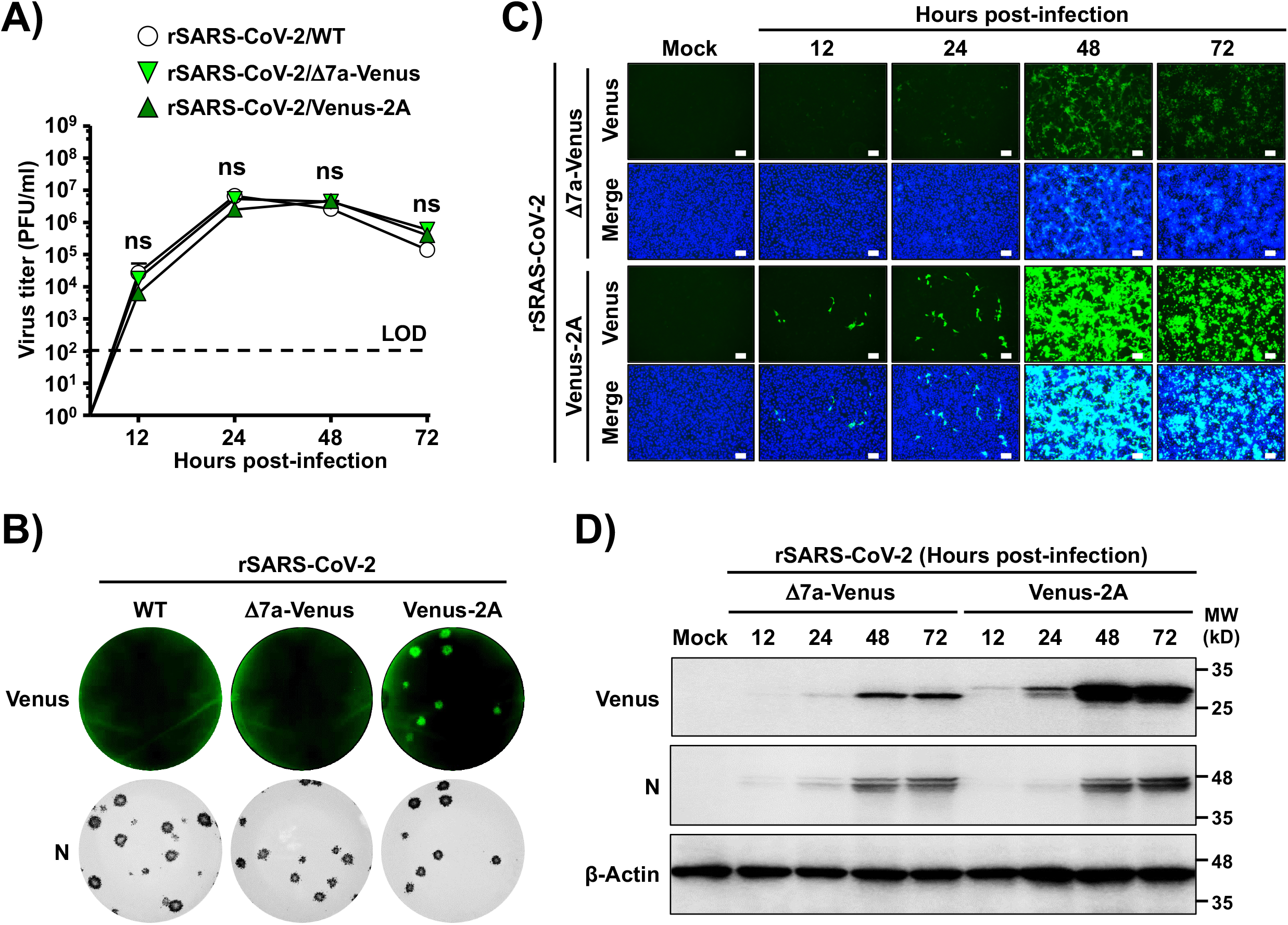
Characterization of rSARS-CoV-2/Venus-2A *in vitro*. **(A)** Tissue culture supernatants from cells infected (MOI 0.01) with rSARS-CoV-2/WT, rSARS-CoV-2/Δ7a-Venus or rSARS-CoV-2/Venus-2A, were collected at the indicated times pi, and viral titers were determined by plaque assay. LOD, limitation of detection; ns, not significant. **(B)** Vero E6 cells infected with ~15 PFU of rSARS-CoV-2/WT (left), rSARS-CoV-2/Δ7a-Venus (middle) or rSARS-CoV-2/Venus-2A (right) were fixed and fluorescent plaques were photographed under a ChemiDoc MP imaging system (top). After imaging, viral plaques were immunostained with the 1C7C7 N protein MAb (bottom). (**C**-**D**) Vero E6 cells infected (MOI 0.001) with rSARS-CoV-2/Δ7a-Venus (top) or rSARS-CoV-2/Venus-2A (bottom) were monitored at the indicated times pi using fluorescent microscopy (**C**). Cell nuclei were stained with DAPI. Scale bars, 100 μm. At the same times pi, whole cell lysate were prepared and analyzed by Western blot analysis using antibodies against Venus and SARS-CoV-2 N protein (1C7C7). β-Actin was used as a loading control (**D**).

### *In vivo* characterization of rSARS-CoV-2/Venus-2A

K18 transgenic mice expressing hACE2 have been shown to be a good animal model of SARS-CoV-2 infection (Oladunni et al., 2020; Zheng et al., 2021). We therefore examined whether SARS-CoV-2 infection could be tracked *ex vivo* using Venus expression. To that end, K18 hACE2 transgenic mice were infected intranasally with 10^5^ PFU of rSARS-CoV-2/Venus-2A, rSARS-CoV-2/Δ7a-Venus or rSARS-CoV-2/WT (**Fig. 3**). Mice were euthanized at 1, 2, 4 and 6 days post-infection (dpi), and their lungs were excised and imaged *ex vivo* using an *in vivo* imaging system (AMI spectrum). Venus expression was readily detected in all lungs obtained from mice infected with rSARS-CoV-2/Venus-2A but not those infected with rSARS-CoV-2/Δ7a-Venus, or rSARS-CoV-2/WT (**Fig. 3A**). Quantitative analyses showed that Venus intensity peaks at 2 dpi and decreases over the course of infection in the lungs of infected mice (**Fig. 3B**). Nevertheless, gross lesions on the lung surface of mice infected with rSARS-CoV-2/Venus-2A was comparable to those observed in rSARS-CoV-2/WT or rSARS-CoV-2/Δ7a-Venus infected mice (**Fig. 3C** and **3D**). In addition, infection with rSARS-CoV-2/Venus-2A and rSARS-CoV-2/Δ7a-Venus resulted in comparable viral titers to those observed in K18 hACE2 transgenic mice infected with rASRS-CoV-2/WT in all organs at all times pi (**Fig. 3E**), suggesting that the undetectable Venus expression in the lungs of K18 hACE2 mice infected with rSARS-CoV-2/Δ7a-Venus is unlikely due to lower levels of viral replication *in vivo*. Notably, we observed a correlation between virus replication and fluorescence intensity in the lungs (**Fig. 3F**).

**Figure 3.**
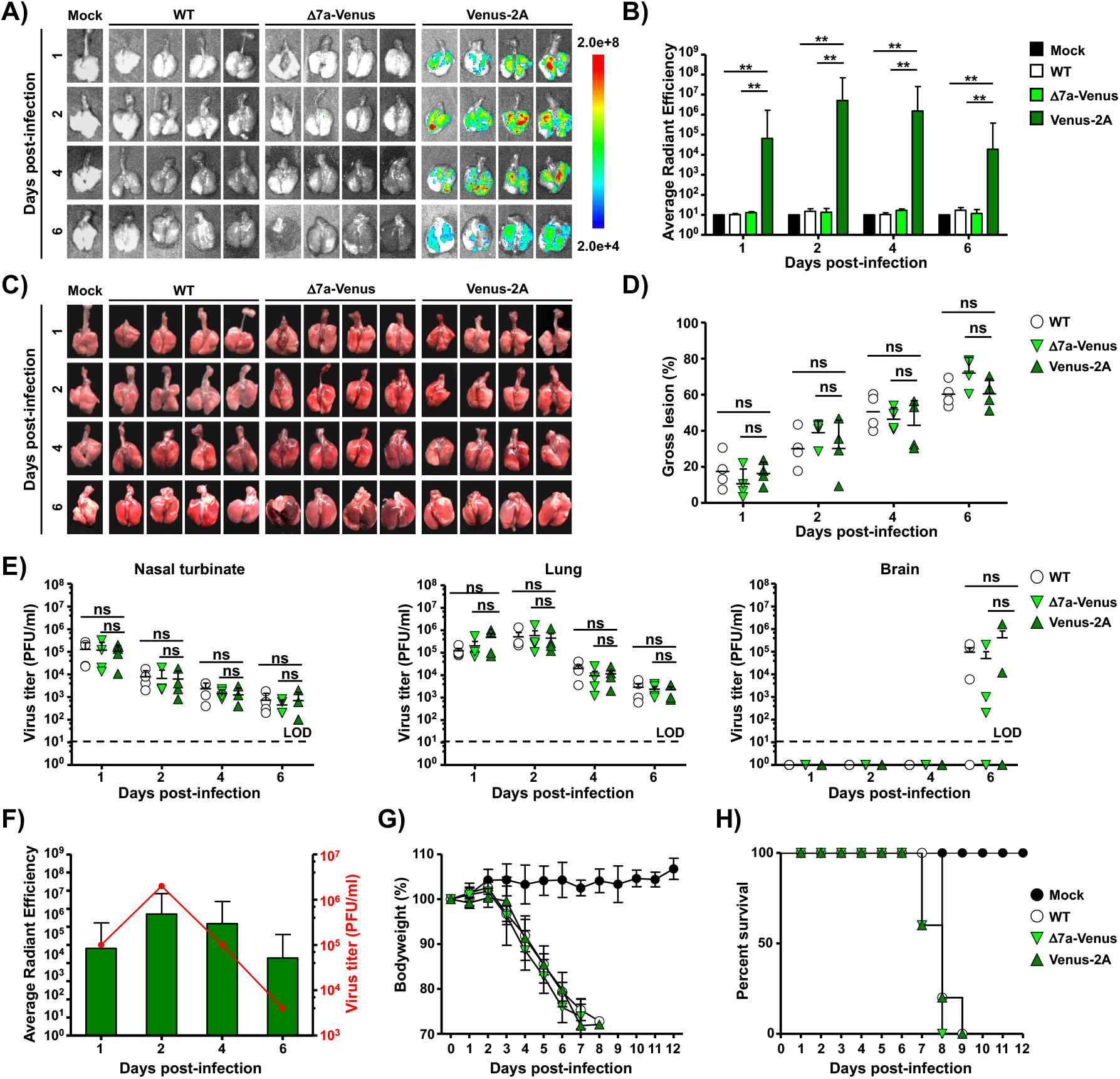
Replication dynamics of rSARS-CoV-2/Venus-2A *in vivo*. (**A-B**) Five-week-old K18 hACE2 transgenic mice were mock-infected or infected (10^5^ PFU/mouse) with rSARS-CoV-2/WT (WT), rSARS-CoV-2/Δ7a-Venus (Δ7a-Venus) or rSARS-CoV-2/Venus-2A (Venus-2A). Lungs were excised at 1, 2, 4 and 6 dpi, and Venus expression was assessed under an IVIS (**A**). Fluorescence intensity was quantitively analyzed by the program of Aura (**B**). (**C**-**D**) Images of lungs were photographed at 1, 2, 4 and 6 dpi (**C**) and the gross lesions on the lung surfaces were quantitively analyzed by ImageJ (**D**). ns, not significant. **(E)** Viral titers in the nasal turbinate (left), lungs (middle) and brain (right) were determined by plaque assay. ns, not significant. **(F)** Correlation between viral titers and Venus intensity in the lungs of rSARS-CoV-2/Venus-2A infected mice. (**G**-**H**) Five-week-old K18 hACE2 transgenic mice were mock-infected or intranasally inoculated with 10^5^ PFU/mouse of rSARS-CoV-2/WT, rSARS-CoV-2/Δ7a-Venus or rSARS-CoV-2/Venus-2A and monitored for 12 days for body weight loss (**G**) and survival (**H**).

To investigate the pathogenicity of rSARS-CoV-2/Venus-2A, we infected (10^5^ PFU) K18 hACE2 transgenic mice intranasally with rSARS-CoV-2/Venus-2A, rSARS-CoV-2/Δ7a-Venus or rSARS-CoV-2/WT, and monitored changes in body weight and survival rate for 12 days after viral infection. All infected mice showed significant bodyweight loss starting from 4 dpi. Mice infected with rSARS-CoV-2/Δ7a-Venus succumbed to viral infection by 8 dpi and mice infected with rSARS-CoV-2/Venus-2A or rSARS-CoV-2/WT succumbed to infection by 9 dpi (**Fig. 3G** and **3H**).

### *In vitro* and *in vivo* stability of rSARS-CoV-2/Venus-2A

Because reporter-expressing rSARS-CoV-2 applications necessitate genetic and phenotypic stability, we evaluated the stability of rSARS-CoV-2/Venus-2A *in vitro* and *in vivo*. To that end, we passaged the rSARS-CoV-2/Venus-2A seven times in Vero E6 cells and tissue culture supernatant (TCS) from selected passages were subjected to the analysis of plaque assay using a fluorescent imaging system and immunostaining with the 1C7C7 N protein monoclonal antibody (MAb) (**Fig. 4A**, left). Viruses present in TCS from P1, P3, P5 and P7 retained 100% Venus expression (**Fig. 4A**, right). For the *in vivo* stability evaluation of rSARS-CoV-2/Venus-2A, lung homogenates from K18 hACE2 transgenic mice at 1, 2, 4 and 6 dpi infected with rSARS-CoV-2/Venus-2A were subjected to plaque assay using a fluorescent imaging system and immunostaining with the 1C7C7 N protein MAb (**Fig. 4B**, left). Lung homogenates from 1 and 2 dpi retained 100% Venus expression, and homogenates from 4 and 6 dpi retained 99% and 98%, respectively, Venus expression (**Fig. 4B**, right).

**Figure 4.**
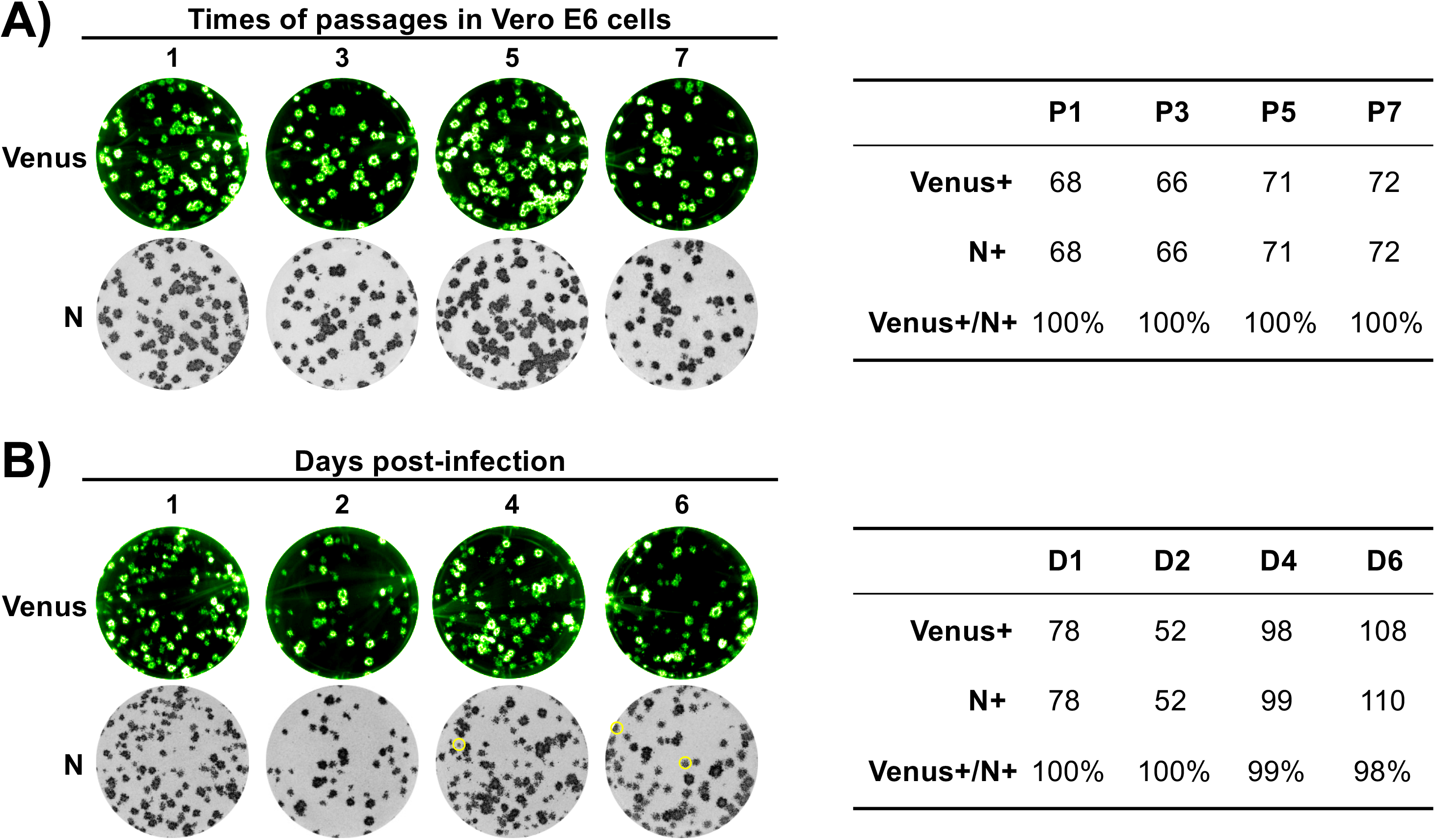
Stability of rSARS-CoV-2/Venus-2A. **(A)** The rSARS-CoV-2/Venus-2A was passaged 7 times in Vero E6 cells and the first (P1), third (P3), fifth (P5) and seventh (P7) passage supernatants were analyzed by plaque assay. Fluorescent plaques were detected under a ChemiDoc MP imaging system (top) and then were immunostained with the 1C7C7 N MAb (bottom). The ratio of Venus positive over N positive plaques was calculated (right). **(B)** The lung from one of the mice described in Figure 3 was homogenized and the clarified supernatant was collected and analyzed by plaque assay. Fluorescent plaques were detected using the ChemiDoc MP imaging system (top) and then immunostained with the N protein MAb 1C7C7 N (bottom). Venus-negative plaques were circled in yellow (bottom). The ratio of Venus-positive over N-positive plaques was calculated (right).

### Visualization of SARS-CoV-2 replication dynamic *in vivo*

While fluorescent Venus expression allowed us to conduct *ex vivo* imaging of lungs from SARS-CoV-2 infected mice, it did not allow us to track viral infection in the entire mouse using IVIS. To circumvent this problem, we engineered a Nluc-expressing rSARS-CoV-2 using our reverse genetic system and the same 2A strategy (**Fig. 5A**). This rSARS-CoV-2/Nluc-2A exhibited a similar plaque phenotype and comparable growth kinetics in Vero E6 cells as the rSARS-CoV-2/WT and our previously described rSARS-CoV-2/Δ7a-Nluc (**Fig. 5B** and **5C**). Notably, Nluc expression levels were increased by more than 30-folds in Vero E6 cells infected with rSARS-CoV-2/Nluc-2A than rSARS-CoV-2/Δ7a-Nluc at 72 hpi (**Fig. 5D**).

**Figure 5.**
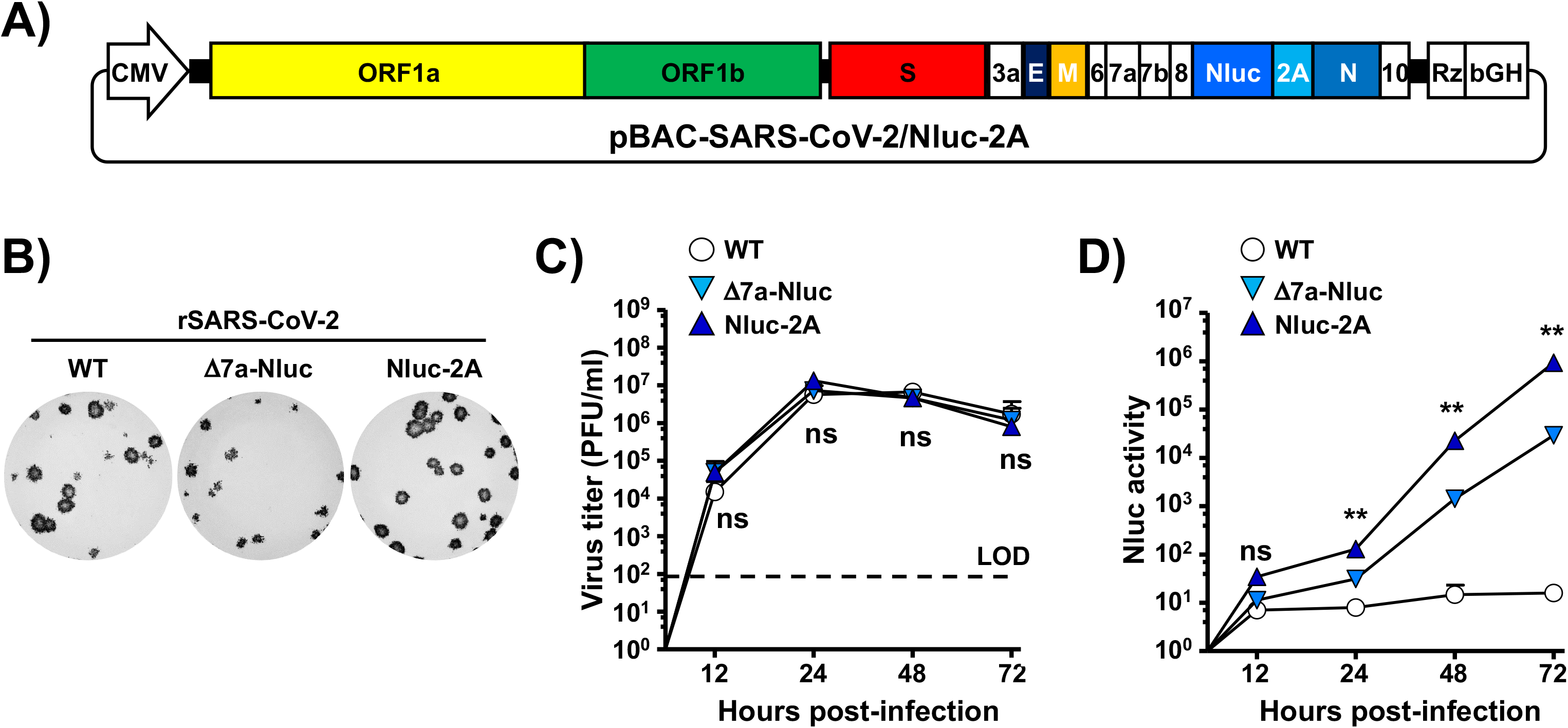
Generation and characterization of rSARS-CoV-2 expressing Nano luciferase-2A *in vitro*. **(A)** Schematic representation of the BAC for generation of rSARS-CoV-2/Nluc-2A. The Nluc coding sequence is indicated by the light blue box. **(B)** Vero E6 cells infected with ~15 PFU of rSARS-CoV-2/WT (left), rSARS-CoV-2/Δ7a-Nluc (middle) or rSARS-CoV-2/Nluc-2A (right) were fixed, permeabilized and immunostained with the 1C7C7 N protein MAb. **(C)** Tissue culture supernatants from Vero E6 cells infected (MOI 0.01) with rSARS-CoV-2/WT, rSARS-CoV-2/Δ7a-Nluc or rSARS-CoV-2/Nluc-2A, were collected at the indicated times pi, and tissue culture supernatants were titrated by plaque assay. LOD, limitation of detection. ns, not significant. **(D)** Tissue culture supernatants from Vero E6 cells infected (MOI 0.01) with rSARS-CoV-2/WT, rSARS-CoV-2/Δ7a-Nluc or rSARS-CoV-2/Nluc-2A, were collected at the indicated times pi and Nluc activity in the tissue culture supernatants was determined. ns, not significant.

Since the rSARS-CoV-2/Nluc-2A expressed significantly higher levels of Nluc than those of our previously described rSARS-CoV-2/Δ7a-Nluc *in vitro*, we evaluated whether SARS-CoV-2 infection could be tracked *in vivo* using Nluc expression directed by rSARS-CoV-2/Nluc-2A. To that end, we infected (10^5^ PFU) K18 hACE2 transgenic mice intranasally with rSARS-CoV-2/Nluc-2A or rSARS-CoV-2/WT (**Fig. 6**). Mice were anesthetized, retro-orbitally injected with Nluc substrate and then imaged under an IVIS at 1, 2, 4 and 6 dpi. Nluc expression was readily detected in mice infected with rSARS-CoV-2/Nluc-2A but not those infected with rSARS-CoV-2/WT (**Fig. 6A**), as previously shown *in vitro* (**Fig. 5D**). Quantitative analyses showed that Nluc intensity continued increasing at later dpi (**Fig. 6B**). Gross lesions on the lung surface of mice infected with rSARS-CoV-2/Nluc-2A was comparable to those in the WT rSARS-CoV-2 infected mice (**Fig. 6C** and **6D**). Importantly, viral titers detected in the rSARS-CoV-2/Nluc-2A infected mice were comparable to those infected with rSARS-CoV-2/WT in all organs tested at different dpi (**Fig. 6E**), despite the Nluc activity was only detected in the organs from rSARS-CoV-2/Nluc-2A virus infected mice (**Fig. 6F**). Unexpectedly, the Nluc intensity did not correlate with the viral titers in the lungs at 4 and 6 dpi (**Fig. 6G**), indicating the Nluc accumulates *in vivo*. Meanwhile, we also compared the pathogenicity of rSARS-CoV-2/Nluc-2A and rSARS-CoV-2/WT in K18 hACE2 transgenic mice. Both rSARS-CoV-2/Nluc-2A and rSARS-CoV-2/WT showed similar virulence (**Figs. 6H** and **6I**).

**Figure 6.**
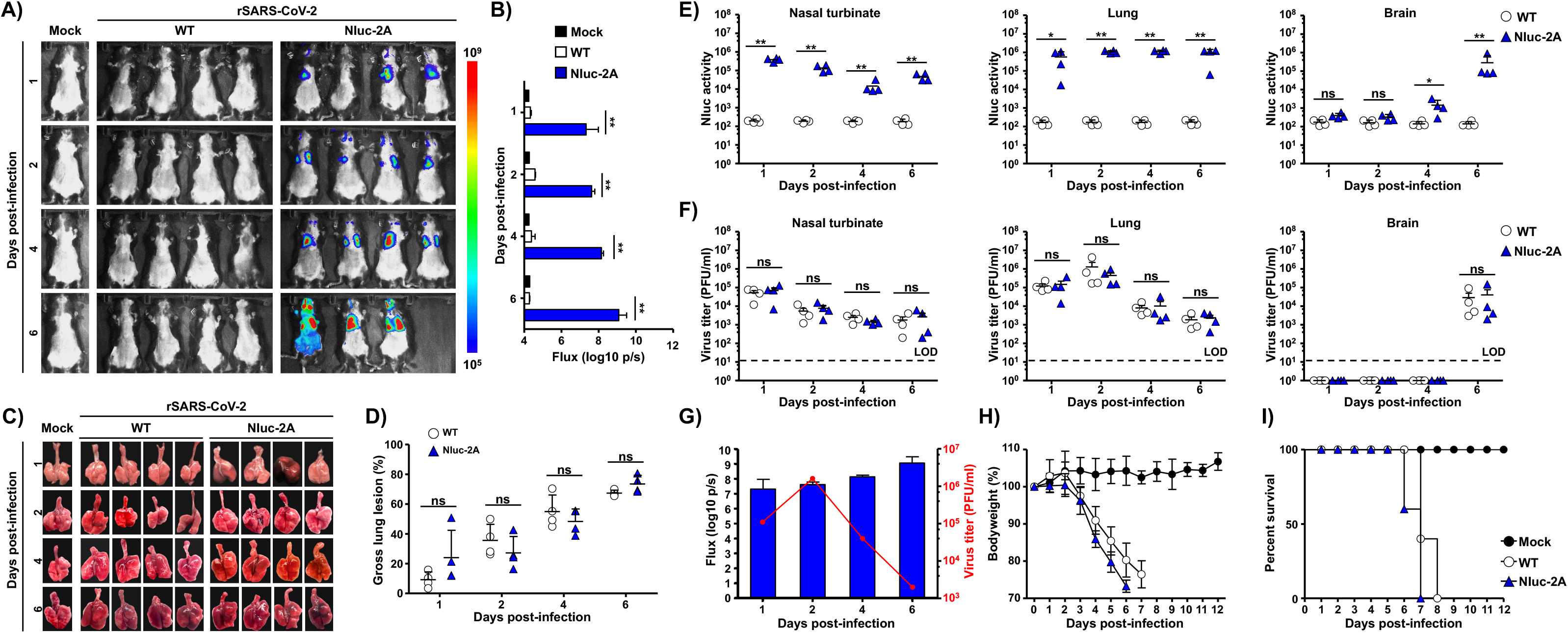
*In vivo* dynamics of SARS-CoV-2 infection by real-time monitoring of Nluc expression. (**A-B**) Five-week-old K18 hACE2 transgenic mice were mock-infected or infected (10^5^ PFU/mouse) with rSARS-CoV-2/WT (WT) or rSARS-CoV-2/Nluc-2A (Nluc-2A). Mice were anesthetized at 1, 2, 4 and 6 dpi and retro-orbitally injected with the Nluc substrate. Nluc expression was determined using an IVIS system (**A**) and quantitively analyzed by the Aura program (**B**). (**C**-**D**) Lungs were excised and photographed at 1, 2, 4 and 6 dpi (**C**), and gross lesions on the lung surfaces were quantitively analyzed by ImageJ (**D**). ns, not significant. **(E)** Nluc activity in the nasal turbinate (left), lungs (middle) and brain (right) from infected mice were determined using a multi-plate reader. ns, not significant. **(F)** Viral titers in the nasal turbinate (left), lungs (middle) and brain (right) were determined by plaque assay. ns, not significant. **(G)** Correlation between viral titers and Nluc intensity in the lungs of rSARS-CoV-2/Nluc-2A-infected mice. (**H**-**I**) Five-week-old K18 hACE2 transgenic mice were mock-infected or infected (10^5^ PFU/mouse) with rSARS-CoV-2/WT or rSARS-CoV-2/Nluc-2A and monitored for 12 days for changes in body weight (**H**) and survival (**I**).

### Effect of NAbs on progression of Nluc-2A virus infection *in vivo*

Although several vaccines against COVID-19 have been already approved for emergency use by United States (US) Food and Drug Administration (FDA), identification and characterization of SARS-CoV-2 NAbs represent a valuable therapeutic option to counteract the putative emergence of variants of concern (VoC). Currently, most NAb screenings are performed in tissue cultured cells rather than *in vivo*, which is heavily reliant upon viral titration of animal organs, a process that is time and labor intensive. We therefore investigated whether the use of rSARS-CoV-2/Nluc-2A could expedite the screening process and facilitate the investigation of how NAb affects the kinetics of virus infection. To that end, we treated K18 hACE2 transgenic prophylactically with 1212C2, a previously described SARS-CoV-2 NAb (Piepenbrink et al., 2021), for 12 h, and then infected them (10^5^ PFU) with rSARS-CoV-2/Nluc-2A. Non-invasive longitudinal imaging of the mice revealed that 1212C2 treated mice dramatically restricted rSARS-CoV-2/Nluc-2A multiplication since no significant Nluc signal was detected at any time point examined (**Figs. 7A** and **7B**). These results were also supported by the observation of reduced lung surface lesions (**Figs. 7C** and **7D**), Nluc expression (**Fig. 7E)** and virus titers **(Fig. 7F)** in the nasal turbinate, lungs, and brains of 1212C2-treated mice. Moreover, 1212C2 was able to protect mice from clinical symptoms of rSARS-CoV-2/Nluc-2A infection as determined by changes in body weight (**Fig. 7G**) and survival rate (**Fig. 7H**).

**Figure 7.**
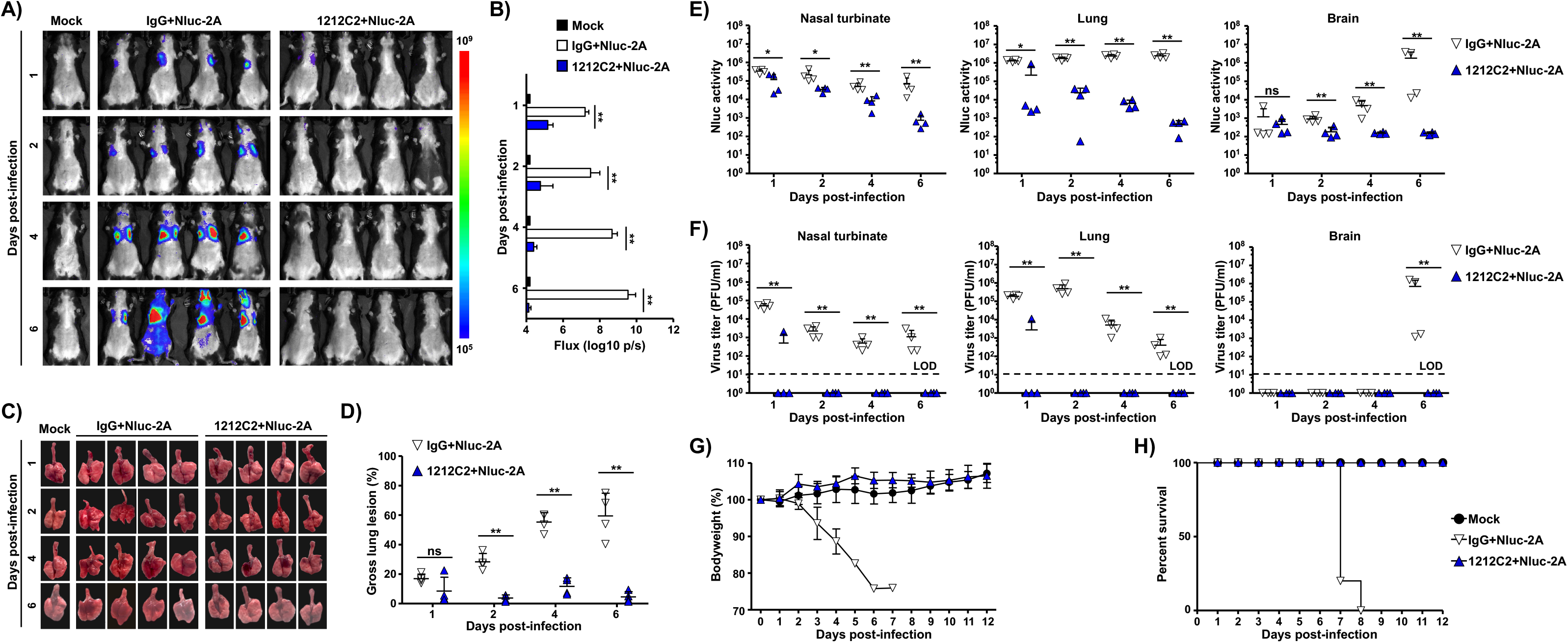
Prophylactic effect of 1212C2 on mice infected with rSARS-CoV-2/Nluc-2A. (**A-B**) Five-week-old K18 hACE2 transgenic mice were injected with isotype IgG control or 1212C2 MAbs and 12 h after treatment, mice were infected (10^5^ PFU/mouse) with rSARS-CoV-2/Nluc-2A (Nluc-2A). Mock-treated and mock-infected mice were included as controls. Mice were anesthetized at 1, 2, 4 and 6 dpi and retro-orbitally injected with the Nluc substrate. Nluc expression was determined using an IVIS system (**A**) and quantitively analyzed by the Aura program (**B**). (**C**-**D**) The lungs were excised and photographed at 1, 2, 4 and 6 dpi (**C**) and the gross lesions on the lung surfaces were quantitively analyzed by ImageJ (**D**). ns, not significant. **(E)** Nluc activity in the nasal turbinate (left), lungs (middle) and brains (right) from infected mice were measured using a multi-plate reader. **(F)** Viral titers in the nasal turbinate (left), lungs (middle) and brain (right) were determined by plaque assay. (**G**-**H**) Five-week-old K18 hACE2 transgenic mice were injected with isotype IgG control or 1212C2 MAbs and 12 h after treatment, mice were infected (10^5^ PFU/mouse) with rSARS-CoV-2/Nluc-2A (Nluc-2A). Mock-treated and mock-infected mice were included as controls. Mice were monitored for 12 days for changes in body weight l (**G**) and survival (**H**).

## DISCUSSION

In this study, we report a novel strategy to generate replication competent reporter-expressing (e.g., Venus or Nluc) rSARS-CoV-2 using the well-documented BAC-based reverse genetic system (Ye et al., 2020a). Our rSARS-CoV-2/Venus-2A and rSARS-CoV-2/Nluc-2A both exhibited rSARS-CoV-2/WT-like growth properties *in vitro* and *in vivo* without displaying attenuation and allowed us to monitor virus infection *ex vivo* in the lungs of infected mice (rSARS-CoV-2/Venus-2A) and the dynamics of viral replication in the entire mouse (rSARS-CoV-2/Nluc-2A) using non-invasive longitudinal *in vivo* imaging. Importantly, we demonstrate the feasibility of using rSARS-CoV-2/Nluc-2A to rapidly identify prophylactics and/or therapeutics *in vivo*.

We and others have previously demonstrated the feasibility of using this 2A approach to generate recombinant viruses expressing reporter genes fused to a viral protein (Caì et al., 2018; Manicassamy et al., 2010; Nogales et al., 2019; Ye et al., 2020b). To generate these novel reporter rSARS-CoV-2 expressing higher level of reporter gene, we placed the PTV-1 2A self-cleaving peptide between the reporter gene and the viral N gene (**Fig. 1**). This results in the expression of a polyprotein that is post-translationally cleaved at the 2A site leading to the individual expression of the reporter gene and the viral N protein (Luke et al., 2010), and the expression of Venus in rSARS-CoV-2/Venus-2A was extremely increased *in vitro* and *in vivo* (**Figs. 2** and **3**, respectively). The rationale of cloning the reporter gene fused to the viral N gene to increase reporter gene expression was based on the N protein being one of the most abundant structural proteins produced during SARS-CoV-2 and other CoVs infections (Hiscox et al., 1995; Hou et al., 2020; Scobey et al., 2013). Importantly, this new 2A approach does not remove any viral gene from the viral genome. Although recent data from our laboratory (Silvas et al., 2021) suggest that ORF7a is not essential for SARS-CoV-2 replication *in vitro* and *in vivo*, it is still largely unknown whether the lack of ORF7a affects some unknown aspects of SARS-CoV-2 infections. In addition, this also increases the instability concern of reporter gene, as shown in a recent study where a reporter gene fused into the C-terminus of ORF7a was not stable (Rihn et al., 2021). Contrarily, our rSARS-CoV-2/Venus-2A exhibited 100% stability after seven passages in cultured cells and retained 98% stability *in vivo* at 6 dpi (**Fig. 4**).

The increased expression level of reporter gene also facilitates the use of bioluminescence imaging of the entire infected mouse. rSARS-CoV-2/Nluc-2A expressed ~30 folds higher levels of Nluc than rSARS-CoV-2Δ7a-Nluc in cultured cells (**Fig. 5**), which allowed us to track the viral infection as early as 1 dpi (**Fig. 6A**). Another advantage is that the rSARS-CoV-2/Nluc-2A could provide a real-time and longitudinal information in a non-invasive manner of infection rather than providing a static “snapshot” using the traditional necropsy and titration of tissues or organs from infected animal. Notably, Nluc activity correlated well with the titers detected in the lungs at 1 and 2 dpi (**Figs. 6E** and **6F**). However, viral titers in lungs were decreased at 4 and 6 dpi, whereas we still detected high levels of Nluc activity. This may reflect Nluc stability (Hall et al., 2012), which may lead to the accumulation and gradual increase of Nluc signal during the course of viral infection. This explanation was also supported by the *in vitro* infection data in which the viral titers were declining yet and the Nluc activity was gradually increasing (**Fig. 5D**).

NAb represent a promising prophylactic and/or therapeutic treatment against SARS-CoV-2, particularly for individuals infected with newly identified VoC. However, evaluation of SARS-CoV-2 NAb *in vivo* relies on the necropsy and viral titration of tissues and/or organs from infected animals. To overcome this limitation, we established a rapid method based on a non-invasive measurement of Nluc expression. By using this *in vivo* imaging system, NAbs could be easily identified as early as 1 dpi and in a relative high throughput method. This was further supported by Nluc activity, viral titration, body weight changes and survival rate (**Fig. 7**), indicating our strategy provides a rapid *in vivo* screening method to identify NAbs against SARS-CoV-2.

In the present work, we have documented a new strategy to generate replication-competent reporter rSARS-CoV-2 expressing higher levels of reporter gene than those previously described by substituting the viral ORF7a protein with the report gene. This novel strategy does not eliminate any viral gene and these new reporter-expressing rSARS-CoV-2 were genetically stable and replicated as efficiently as rSARS-CoV-2/WT both *in vitro* and *in vivo*, with comparable pathogenicity in K18 hACE2 transgenic mice. Notably, the robust levels of reporter gene expression of these new reporter-expressing rSARS-CoV-2 represent an excellent option to study viral pathogenesis, tissue tropism, and replication kinetics of SARS-CoV-2, including recently identified VoC.

## MATERIAL AND METHODS

### Biosafety

All the *in vitro* and *in vivo* experiments with infectious rSARS-CoV-2 were conducted under appropriate biosafety level (BSL) 3 and animal BSL3 (ABSL3) laboratories, respectively, at Texas Biomedical Research Institute (Texas Biomed). Experiments were approved by the Texas Biomed Institutional Biosafety (IBC) and Animal Care and Use (IACUC) committees.

### Cells and viruses

African green monkey kidney epithelial cells (Vero E6, CRL-1586) were obtained from the American Type Culture Collection (ATCC, Bethesda, MD) and maintained in Dulbecco’s modified Eagle medium (DMEM) supplemented with 5% (v/v) fetal bovine serum (FBS, VWR) and 1% penicillin-streptomycin-glutamine (PSG) solution (Corning).

Recombinant (r)SARS-CoV-2 were generated based on the backbone of the USA-WA1/2020 strain using a previously described bacterial artificial chromosome (BAC)-based reverse genetics system (Chiem et al., 2020; Ye et al., 2020a).

### Rescue of rSARS-CoV-2

Virus rescue experiments were performed as previously described (Avila-Perez et al., 2019). Briefly, confluent monolayers of Vero E6 cells (10^6^ cells/well, 6-well plates, triplicates) were transfected with 4.0 μg/well of SARS-CoV-2 BAC using Lipofectamine 2000. After 24 h, transfection media was exchanged for post-infection media (DMEM supplemented with 2% FBS and 1% PSG), and cells were split and seeded into T75 flasks 72 h post-transfection. After incubation for another 72 h, tissue culture supernatants were collected, labeled as P0 and stored at −80°C. After being titrated, the P0 virus was used to infect fresh Vero E6 cells at MOI 0.0001 for 72 h to generate P1 stocks. P1 viral stocks were aliquoted and stored at −80°C until being used.

### Western blot

Whole cell lysates, SDS-PAGE, and Western blotting were performed as previously described (Ye et al., 2020b). Briefly, cells were lysed in passive lysis buffer (Promega, MI, USA) at 4°C for 30 min, followed by centrifugation at 12,000 g at 4°C for another 30 min. Equivalent amounts of cell lysates were subjected to 12% SDS-PAGE and transferred to nitrocellulose membranes. After blocking with 5% bovine serum albumin in PBS containing 0.1% Tween 20 at room temperature for 1 h, the membranes were incubated with the indicated primary antibodies at 4°C overnight, followed by horseradish peroxidase-conjugated secondary antibody incubation at 37°C for 1 h. β-Actin was used as the loading control. Membranes were developed with ECL detection reagent (Thermo Fisher Scientific) in the ChemiDoc MP Imaging System (Bio-Rad, Hercules, CA, USA).

### Immunofluorescence assay (IFA)

Vero E6 cells (10^6^ cells/well, 6-well plate format, triplicates) were mock-inoculated or inoculated with the tissue culture supernatant (1 ml/well) of Vero E6 cells transfected with the BACs collected at 72 h post-transfection. At 48 h post-inoculation, cells were fixed with 10% formaldehyde solution at 4°C overnight and permeabilized using 0.5% (v/v) Triton X-100 in PBS for 15 min at room temperature. Then, cells were incubated overnight with 1 μg/ml of a SARS-CoV cross-reactive N MAb (1C7C7) at 4°C, washed stringently with PBS, and stained with a FITC-labeled goat anti-mouse IgG (1:200). Finally, cells were visualized and imaged under a EVOS fluorescent microscope (Thermo Fisher Scientific).

### RT-PCR

Total RNA from virus infected (MOI=0.01) Vero E6 cells (10^6^ cells/well, 6-well plate format) was extracted with TRIzol Reagent (Thermo Fisher Scientific) according to the manufacturer’s instructions. RT-PCR amplification of the viral genome spanning nucleotides 27,895-29,534 (according to the SARS-CoV-2 USA-WA1/2020 viral genome) was performed using Super Script II Reverse transcriptase (Thermo Fisher Scientific) and Expanded High Fidelity PCR System (Sigma Aldrich). The amplified DNA products were separated on a 0.7% agarose gel. All primer sequences used for RT-PCR are available upon request.

### Plaque assay and immunostaining

Confluent monolayers of Vero E6 cells (10^6^ cells/well, 6-well plate format, triplicates) were infected with serial viral dilutions for 1 h at 37°C. After viral adsorption, cells were overlaid with pi media containing 1% low melting agar and incubated at 37°C. At 72 hpi, cells were fixed overnight with 10% formaldehyde solution. For visualization of Venus, plates were photographed under a ChemiDoc MP Imaging System. For immunostaining, cells were permeabilized with 0.5% (v/v) Triton X-100 in PBS for 15 min at room temperature and immunostained using the SARS-CoV cross-reactive N protein 1C7C7 MAb (1 μg/ml) and the Vectastain ABC kit (Vector Laboratories), following the manufacturers’ instruction. After immunostaining, plates were scanned and photographed using a ChemiDoc MP Imaging System.

### Virus growth kinetics

Confluent monolayers of Vero E6 cells (6-well format, 10^6^ cells/well, triplicates) were mock infected or infected (MOI=0.01) with rSARS-CoV-2/WT, rSARS-CoV-2/Δ7a-Venus, rSARS-CoV-2/Venus-2A or rSARS-CoV-2/Nluc-2A. After 1 h of virus adsorption at 37°C, cells were washed with chilled PBS and overlaid with 3 ml of pi medium and incubated at 37°C. At the indicated times pi (12, 24, 48 and 72 h), viral titers in the tissue culture supernatants were determined by plaque assay (Nogales et al., 2014). Presence of Nluc in the tissue culture supernatants from mock and rSARS-CoV-2/WT, rSARS-CoV-2/Δ7a-Nluc or rSARS-CoV-2/Nluc-2A infected cells was quantified using Nano-Glo® Luciferase Assay System (Promega) following the manufacturers’ specification.

### Animal experiments

All animal protocols were approved by Texas Biomed IACUC (1718MU). Five-week-old female K18 hACE2 transgenic mice were purchased from The Jackson Laboratory and maintained in the animal facility at Texas Biomed under specific pathogen-free conditions. For virus infection, mice were anesthetized following gaseous sedation in an isoflurane chamber and inoculated intranasally with a dose of 10^5^ PFU/mouse.

For *ex vivo* imaging of lungs, mice were humanely euthanized at 1, 2, 4 and 6 dpi to collect lungs. Fluorescent images of lungs were photographed using an *in vivo* imaging system (AMI HTX) and the bright field images of lungs were taken using an iPhone 6s (Apple, CA, USA). Nasal turbinate, lungs and brains from mock or infected animals were homogenized in 1 ml of PBS for 20 s at 7,000 rpm using a Precellys tissue homogenizer (Bertin Instruments, Yvelines, France). Tissue homogenates were centrifuged at 12,000 g (4 °C) for 5 min, and supernatants were collected and titrated by plaque assay and immunostaining as previously described.

For *in vivo* bioluminescence imaging, mice were anesthetized with isoflurane, injected retro-orbitally with 100 μl of Nano-Glo luciferase substrate (Promega), and immediately imaged. The bioluminescence data acquisition and analysis were performed using the Aura program (AMI spectrum). Flux measurements were acquired from the region of interest. The scale used is included in each figure. Immediately after imaging, nasal turbinate, lungs, and brains were collected and homogenized in 1 ml of PBS. Supernatants were collected and presence of virus was determined by plaque assay and immunostaining, as described above. Nluc activity in the tissue culture supernatants was determined under a multiplate reader (BioTek Instruments, Inc) as above.

For the body weight and survival studies, five-week-old female K18 hACE2 transgenic mice were infected intranasally with 10^5^ PFU/animal following gaseous sedation in an isoflurane chamber. After infection, mice were monitored daily for morbidity (body weight) and mortality (survival rate) for 12 days. Mice showing a loss of more than 25% of their initial body weight were defined as reaching the experimental end-point and humanely euthanized, and the survival curves were plotted according to the method of Kaplan-Meier (Efron, 1988).

### Statistical analysis

All data is presented as mean ± standard deviation (SD) for each group and analyzed by SPSS13.0 (IBM, Armonk, NY, USA). A *P* value of less than 0.05 (*P*<0.05) was considered statistically significant.

## ACKNOWLEDGEMENTS

We are grateful to Dr. Thomas Moran at The Icahn School of Medicine at Mount Sinai for providing the SARS-CoV cross-reactive 1C7C7 N protein MAb.

## AUTHOR CONTRIBUTIONS

C.Y rescued the recombinant viruses and conducted the *in vitro* characterization; C.Y, K.C, J.P, J.A.S and D.M.V conducted the *in vivo* experiments; J.B.T, J.J.K and M.R.W provided critical reagents; J.C.T deep sequenced the viruses; C.Y drafted the manuscript; J.B.T, J.J.K, M.R.W and J.C.T revised the manuscript; L.M.S conceived the study, provided funding, revised and finalized the manuscript.

